# A four-dimensional single-cell and spatial atlas of cAMP-driven repair after rat spinal cord injury

**DOI:** 10.64898/2026.11.15.766731

**Authors:** Quique Vidal Beneyto, Loris Mannino, Guillem Paniagua Soriano, Eric López-Mocholi, Andrés Colomer, Sara Torres Rodríguez, Arnau Montagud, Sonia Tarazona, Victoria Moreno Manzano, Ana Conesa

**Affiliations:** PhD Program in Biotechnology, Universitat Politècnica de València (UPV), València, Spain; Genomics of Gene Expression Lab, Institute for Integrative Systems Biology (I2SysBio), Consejo Superior de Investigaciones Científicas (CSIC)–Universitat de València (UV), Paterna, Spain; Departamento de Estadística e Investigación Operativa Aplicadas y Calidad, Universitat Politècnica de València (UPV), València, Spain; Neuronal and Tissue Regeneration Lab, Centro de Investigación Príncipe Felipe (CIPF), València, Spain; ⁵Systems Biology Modelling Lab, Institute for Integrative Systems Biology (I2SysBio), Consejo Superior de Investigaciones Científicas (CSIC)–Universitat de València (UV), Paterna, Spain

## Abstract

Cyclic adenosine monophosphate (cAMP) induction through photoactivatable adenylate cyclase (bPAC) in corticospinal neurons enhances locomotion after spinal cord injury (SCI) [1]; however, the molecular and cellular mechanisms that underlie this improvement are unknown. Here we perform single-nucleus RNA sequencing (snRNA-seq) and spatial transcriptomics (ST) of spinal cord sections at the injury site of dorsal hemisected rats, to identify the pathways through which bPAC stimulation exerts its regenerative cues after SCI, along with the changes in cellular populations that occur in different parts of the spinal cord.

## Introduction

Upon diverse forms of trauma at the thoracic level of the spinal cord, such as contusion or hemisection, voluntary motor control is suppressed, usually irreversibly. This injury is known as spinal cord injury (SCI), and its physiological explanation is the disruption of connectivity between supraspinal inputs and lower spinal neurons, which renders them non-functional.

In recent years, work using an optogenetic bPAC device has demonstrated the therapeutic potential of cAMP induction in the corticospinal tract (CST) for restoring voluntary motor control after SCI [1]. However, the molecular pathways by which cAMP mediates this motor recovery have remained largely unknown. In this study, we present a comprehensive experimental design in which we explore the cellular and molecular consequences of bPAC optogenetic stimulation in injured rats, using single-nucleus RNA sequencing and spatial transcriptomics.

In our study, single-nucleus RNA-seq serves a twofold purpose. On one hand, we construct a map of the cell types present in uninjured, injured and stimulated spinal cords, to better understand the differences at cell-type level between experimental conditions. On the other hand, it provides a precise reference for spatial transcriptomics mapping: while single-nucleus RNA sequencing provides exceptional cellular granularity, it lacks the spatial context required to determine how cell types are organized relative to the injury epicenter. To map these spatial dynamics, Visium spatial transcriptomics [20] was employed to resolve region-specific cellular remodeling.

Therefore, to characterize the cell-type-specific and spatially resolved responses of the spinal cord to bPAC optogenetic stimulation following SCI, we performed snRNA-seq paired with ST across five experimental conditions: uninjured controls (SHAM), injured animals at 10 and 30 days post-injury (CTRL10, CTRL30), and injured animals receiving bPAC stimulation at 10 and 30 days (STIM10, STIM30). SnRNA-seq samples were obtained rostral to the injury site, whereas ST samples were obtained at both rostral and caudal sections (Fig. 1A).

**Figure 1.**
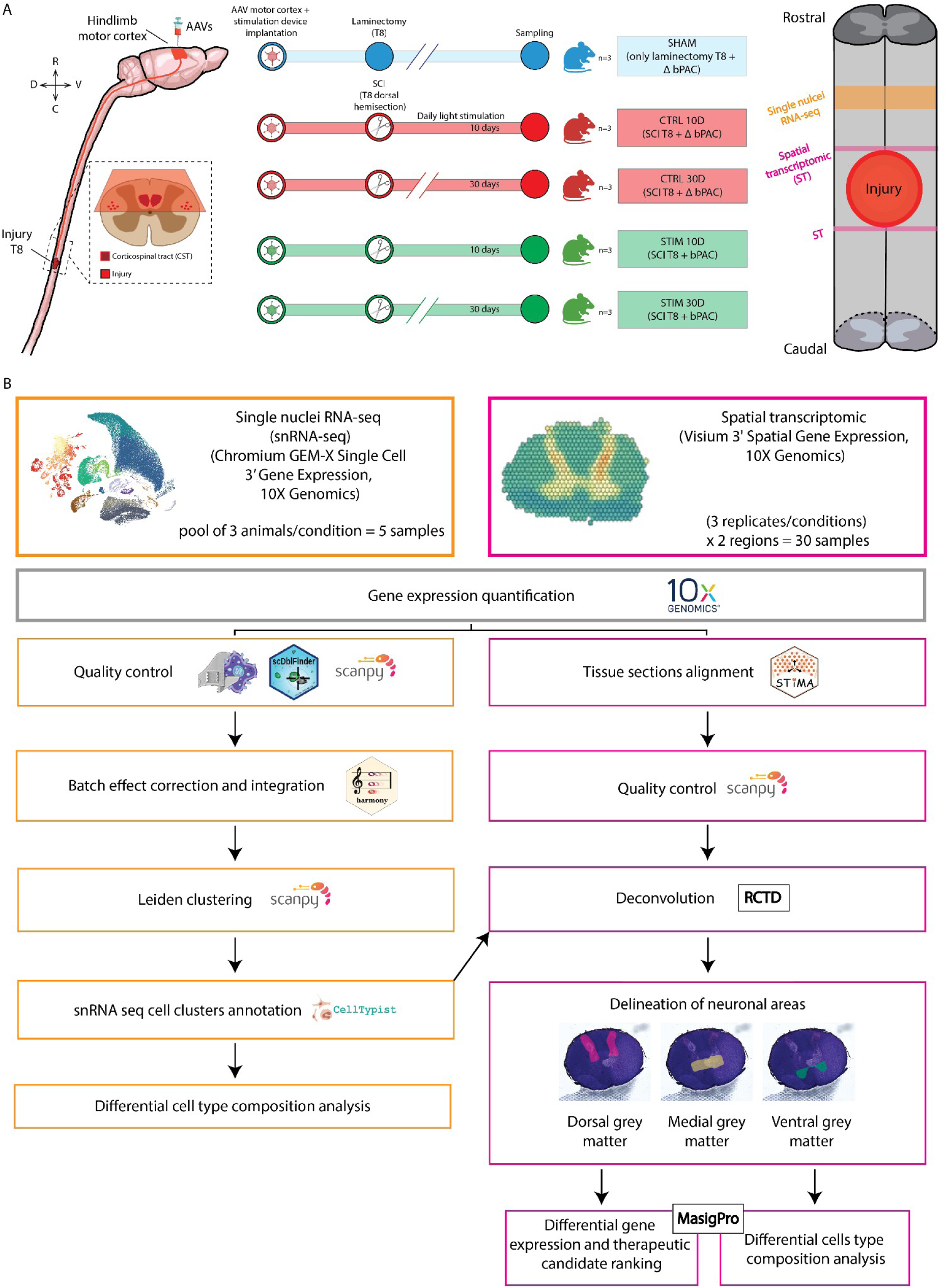
Experimental design and pipeline of bioinformatic analysis. **A,** Rats received AAVs carrying bPAC, or the inactive mutant ΔbPAC as a control, in the hindlimb motor cortex, and the light-stimulation device was implanted at the same time. There were five groups (n = 3 each): • SHAM: T8 laminectomy + ΔbPAC • CTRL: T8 dorsal hemisection + ΔbPAC, at 10 or 30 days • STIM: T8 dorsal hemisection + bPAC, at 10 or 30 days The SCI groups received daily light stimulation until sampling. Tissue rostral to the injury was used for snRNA-seq (orange). Sections rostral and caudal to the lesion were used for spatial transcriptomics (ST, magenta). **B,** Pipeline of bioinformatics analysis: • **snRNA-seq** (3 rats pooled per condition, n = 5): quality control, Harmony integration, Leiden clustering, CellTypist annotation and differential cell-type composition analysis. • **Visium ST** (3 replicates × 5 conditions × 2 regions, n = 30): section alignment (STIMA), quality control and RCTD deconvolution using the snRNA-seq reference. Dorsal, medial and ventral grey matter were then delineated for differential expression (maSigPro), candidate ranking and cell-type composition analysis.

## Results

### A four-dimensional single-cell and spatial atlas of cAMP-driven spinal cord repair

To study the cellular response of the spinal cord to cAMP-dependent stimulation of the corticospinal tract, using an optogenetic approach based on the bacterial photoactivated adenylyl cyclase bPAC [2], after thoracic spinal cord injury, we designed a replicated multi-platform intervention study that resolves recovery in four dimensions at cellular resolution. Spatial transcriptomics captured the two dimensions of each tissue section; sampling immediately both rostral and caudal to the lesion added the rostro-caudal axis as a third dimension; and profiling at two post-injury timepoints added time as a fourth. The design comprised five experimental conditions — uninjured animals (SHAM), injured animals at 10 and 30 days post-injury (CTRL10, CTRL30), and injured animals receiving bPAC stimulation at the same two timepoints (STIM10, STIM30) — each with three biological replicates. For the spatial arm, this yielded 30 samples (3 replicates × 5 conditions × 2 areas). Single-nucleus RNA sequencing was performed on the same animals, pooled by experimental condition, giving five snRNA-seq samples from contiguous rostral segments (Fig. 1A). Deconvolution of each spatial sample drew on the snRNA-seq reference matched to its own experimental condition, ensuring that condition-specific cell types were represented in the reference rather than smoothed away by a single pooled atlas. To our knowledge, this is the first four-dimensional dataset of spinal cord injury recovery, and the most comprehensive assembled to date.

Bioinformatic analysis combined well-established methods with two strategies developed for the specific demands of this design (Fig. 1B). Spinal cord slices were aligned with STIMA [3], so that equivalent anatomical regions were compared across replicates and conditions rather than across arbitrary section coordinates. Temporal dynamics were modelled with maSigPro [4,5] in three different scenarios. First, this allowed us to resolve how gene expression changes over time and how those trajectories diverge between bPAC-stimulated and unstimulated animals in ST samples. Second, it allowed us to resolve how cell-type abundance and cell-type colocalization change over time and between conditions, helping us map the changes in local neighbourhoods of the spinal cord tissue.

### bPAC stimulation preserves global cell-type composition while remodelling ventral and progenitor neuronal populations

Single-nucleus profiling resolved the major cell types of the spinal cord (Fig. 2A) and, within the neuronal compartment, their subtypes (Fig. 2B). The atlas comprised the full oligodendrocyte lineage (OPC, NFOL, MFOL, MOL), astrocytes, ependymal cells, homeostatic and reactive microglia, pericytes and vascular endothelial cells, peripheral immune cells, and six broad neuronal classes (dorsal excitatory, DE; dorsal inhibitory, DI; medial excitatory, ME; medial inhibitory, MI; ventral excitatory, VE; ventral inhibitory, VI). Annotations, based on reported mouse datasets [6–17], were refined manually and validated against canonical markers, which showed the expected cluster-restricted expression (Pdgfra and Cspg4 for OPCs, Tmem119 and P2ry12 for microglia, Gja1 and Aqp4 for astrocytes, Snap25, Slc17a6 and Gad1 for neuronal populations; Fig. 2C, D).

**Figure 2.**
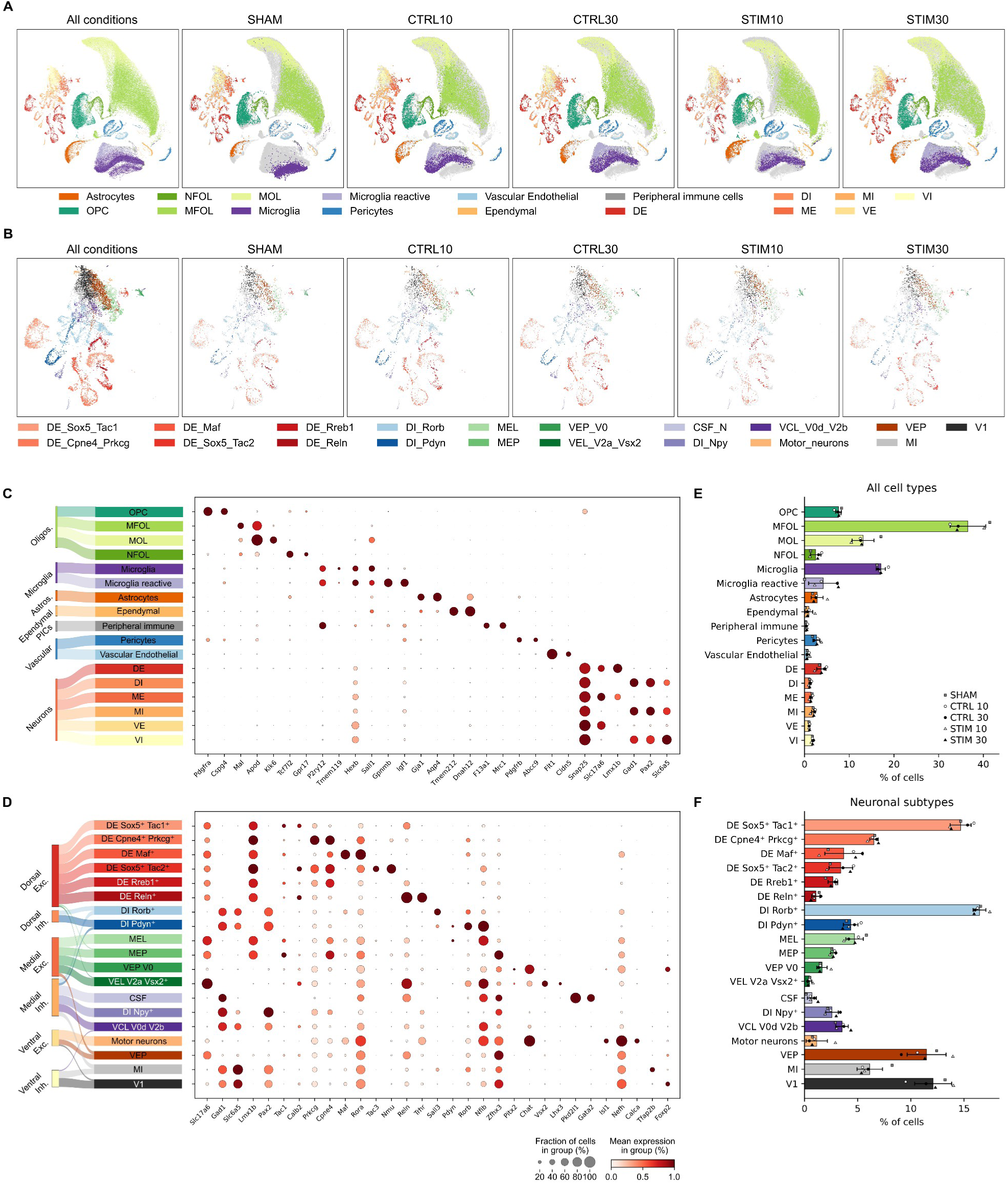
Cellular and neuronal diversity in the spinal cord revealed by single-nucleus RNA sequencing. **A,** UMAP embedding of all cell types identified in the snRNA-seq dataset, shown for all conditions combined and split by condition (SHAM, CTRL10, CTRL30, STIM10, STIM30). Cells are coloured by main cell type identity: oligodendrocyte lineage (OPC, NFOL, MFOL, MOL), astrocytes, ependymal cells, homeostatic and reactive microglia, pericytes and vascular endothelial cells, peripheral immune cells, and the six major neuronal classes (DE, DI, ME, MI, VE, VI). Cells of other conditions are shown in grey in each split panel. **B,** UMAP embedding restricted to neuronal populations, subclustered into transcriptionally distinct subtypes and shown for all conditions combined and split by condition, coloured according to the legend. **C,** Left, flow diagram linking the broad compartments (oligodendrocytes, microglia, astrocytes, ependymal cells, peripheral immune cells, vascular cells and neurons) to the annotated main cell types. Right, dot plot of the canonical marker genes used to annotate each main cell type. Dot size represents the fraction of cells in the group expressing the gene and colour intensity the scaled mean expression, as indicated in the shared key. **D,** As in C, for the neuronal subtypes, grouped by dorsal excitatory, dorsal inhibitory, medial excitatory, medial inhibitory, ventral excitatory and ventral inhibitory classes, with the literature-derived markers used for their annotation. **E,** Relative abundance of each main cell type, as percentage of cells. Bars give the mean across all samples, with individual samples overlaid: SHAM (square), CTRL10 (open circle), CTRL30 (filled circle), STIM10 (open triangle), STIM30 (filled triangle). **F,** As in E, for the neuronal subtypes.

Global composition was preserved across the five conditions: every major cell type was detected at comparable representation in SHAM, injured and stimulated animals (Fig. 2A, E). Oligodendrocyte-lineage cells dominated the dataset, followed by the microglial populations, as expected (Fig. 2E). The orderly segregation of both oligodendrocyte and microglial clusters reproduces the lineage structure of the uninjured and injured reference atlases, and confirms the annotation of the compartment that underlies demyelination and remyelination after SCI, represented by the CTRL and STIM groups (Fig. 2E, F). These cell population proportions match those expected from nucleus-dissociation protocols applied to whole tissue blocks of the spinal cord [18].

To resolve neuronal heterogeneity further, we subclustered the neuronal compartment in our data and in the reference atlas and applied the same CellTypist-based annotation [19], yielding 19 transcriptionally distinct subtypes: six dorsal excitatory (DE Sox5⁺Tac1⁺, DE Cpne4⁺Prkcg⁺, DE Maf⁺, DE Sox5⁺Tac2⁺, DE Rreb1⁺, DE Reln⁺), three dorsal inhibitory (DI Rorb⁺, DI Pdyn⁺, DI Npy⁺), a set of medial and ventral excitatory and inhibitory populations (MEL, MEP, VEP V0, VEL V2a Vsx2⁺, VCL V0d V2b, VEP, MI), cerebrospinal fluid-contacting (CSF) neurons and V1 interneurons. Initially, we also detected neural stem cells, but we considered them a label-transfer artefact and removed them (see Methods). We also manually added motor neurons, as they were absent from the reference dataset (Fig. 2B). Subtype markers behaved as reported in the reference atlas [17], with graded, subtype-restricted expression across clusters (Sox5 and Tac1 for DE Sox5⁺Tac1⁺ neurons, Chat for motor neurons, Pkd2l1 for CSF neurons; Fig. 2D).

Taken together, these findings indicate that injury and stimulation largely preserve the broad neuronal class composition of the spinal cord, comparable to cell profiles previously described in mice [6,7,10,11]. However, due to the unreplicated nature of the single-nucleus dataset, we avoided comparisons across conditions and drew no conclusions about possible cellular changes from this arm alone.

### Spatially resolved deconvolution separates general injury responses from stimulation-specific remodelling

To determine how injury and cortical cAMP stimulation influence the cellular composition of the spinal cord, and how that composition is spatially reorganized, we profiled the sections immediately rostral and caudal to the injury core from all five conditions using Visium spatial transcriptomics [20]. This technology profiles the transcriptome of 55 µm spots on the tissue and is therefore not suitable for resolving individual cells, as each capture area is large enough to contain more than one cell. We therefore used Robust Cell Type Decomposition (RCTD) [21] to deconvolve the cell-type composition of every spot, using our snRNA-seq dataset as reference (Fig. 3A). The deconvolved distributions recovered the expected cell distribution along spinal cord anatomy, with mature oligodendrocyte-lineage and glial populations enriched in white matter, whereas neuronal populations concentrated in grey matter, in every condition and at both rostro-caudal levels.

**Figure 3.**
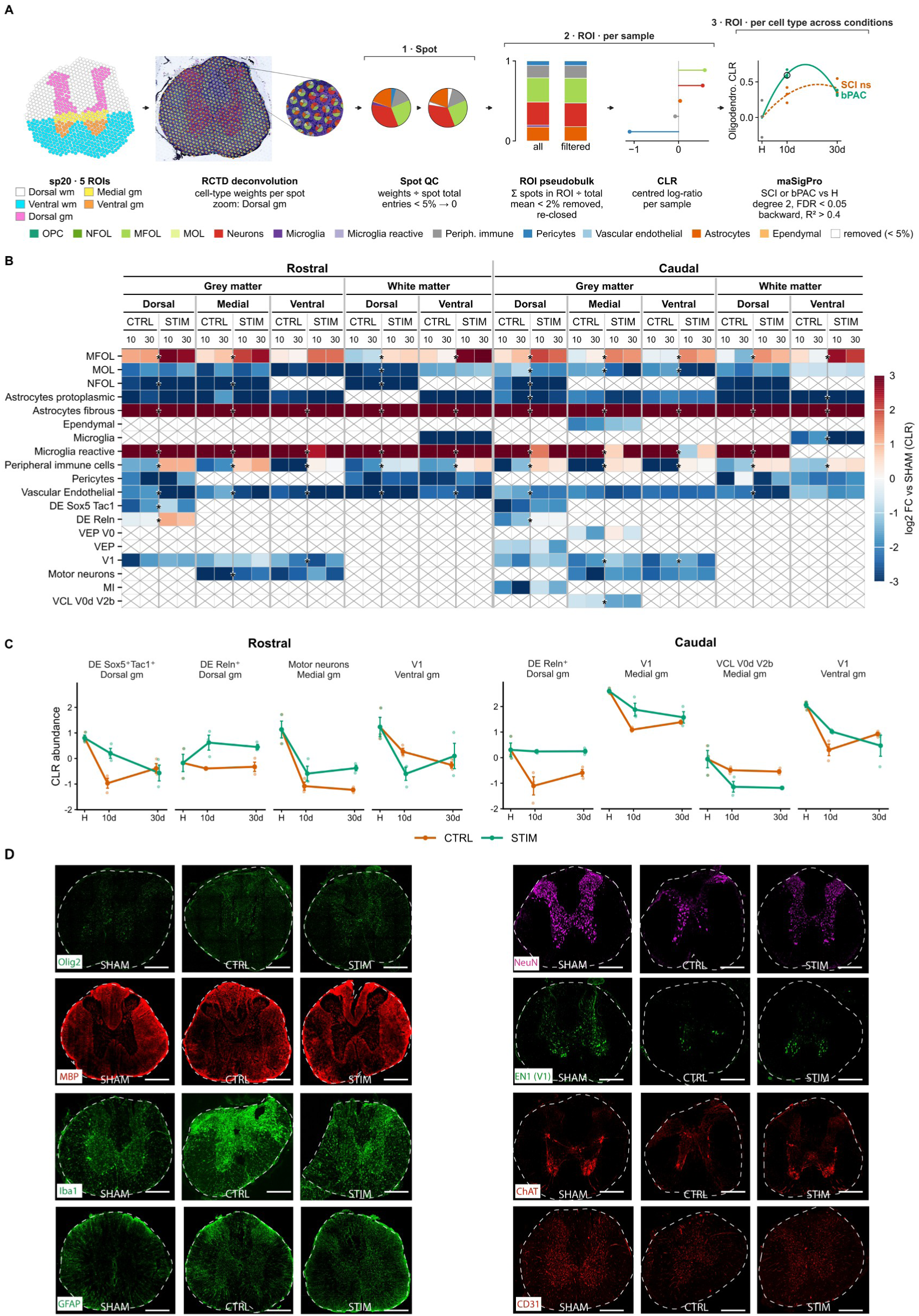
Spatial transcriptomic mapping of cell type and neuronal subtype distribution across spinal cord ROIs following bPAC-mediated optogenetic stimulation. **A,** Analysis pipeline. Left, the five ROIs delineated on each section (dorsal and ventral white matter; dorsal, medial and ventral grey matter), shown for a representative sample, and the RCTD deconvolution returning cell-type weights per spot, with a zoom on dorsal grey matter showing per-spot composition. Step 1, spot level: weights divided by the spot total, entries below 5% set to zero. Step 2, ROI level per sample: spot weights summed within each ROI and divided by the total, cell types below 2% mean abundance removed and compositions re-closed, then centred log-ratio (CLR) transformed per sample. Step 3, ROI level per cell type across conditions: CLR abundances modelled with maSigPro (degree 2, FDR < 0.05, backward selection, R² > 0.4), comparing the SCI and bPAC trajectories against the SHAM baseline. **B,** Heatmap of log2 fold-change in CLR-normalized abundance relative to SHAM, for main cell types and neuronal subtypes, stratified by segment (rostral, caudal), tissue compartment (grey matter: dorsal, medial, ventral; white matter: dorsal, ventral), condition (CTRL, STIM) and timepoint (10, 30 days). Crossed cells indicate a cell type that did not meet the 2% mean abundance threshold in that ROI. Asterisks denote a significant STIM versus CTRL trajectory identified by maSigPro. Colour scale from −3 (blue, decreased) to +3 (red, increased). **C,** CLR abundance trajectories across SHAM (H), 10 and 30 days for the neuronal subtypes whose trajectory differed significantly between CTRL and STIM, in rostral (left) and caudal (right) segments. Data are mean ± SEM with individual replicates overlaid; CTRL in orange, STIM in green. **D,** Representative immunofluorescence of coronal spinal cord sections from SHAM, CTRL and STIM animals at 10 dpi. Left column: Olig2 (green), MBP (red), Iba1 (green) and GFAP (green). Right column: NeuN (magenta), EN1 marking V1 interneurons (green), ChAT (red) and CD31 (red). Dashed lines outline the section border; scale bars as indicated.

The analytical pipeline proceeded through three levels, from the single spatial barcode to the condition-level comparison (Fig. 3A). First, at the spot level, the spatial barcodes of each section were demarcated into five regions of interest (ROIs: dorsal white matter, ventral white matter, and dorsal, medial and ventral grey matter) guided by H&E morphology, and the cell-type proportions within each spot were obtained using RCTD [21] trained on our matched, marker-validated snRNA-seq reference (Fig. 2). Spot-level weights were then quality-controlled, with weights divided by the spot total and entries below 5% set to zero. Second, at the ROI level per sample, the surviving spot weights were aggregated into ROI pseudobulk compositions: the sum of spot weights within each ROI divided by the total, with cell types below 2% mean abundance removed and compositions re-closed, then transformed by the centred log-ratio (CLR) per sample. Third, at the ROI level per cell type across conditions, the CLR-transformed abundances of each cell type were modelled across the time course (SHAM, 10 days, 30 days) with maSigPro [4,5] (degree 2, FDR < 0.05, backward selection, R² > 0.4), comparing the injured and bPAC-stimulated trajectories against the SHAM baseline, separately for each ROI and for each axis, rostral and caudal.

Comparing ROI-specific cell-type abundance against SHAM (Fig. 3B) separated stimulation-independent injury responses from stimulation-specific ones. Fibrous astrocytes and reactive microglia were scarcely represented in SHAM samples and were markedly increased in injured animals, both CTRL and STIM. In stimulated animals at 10 days after cAMP delivery, reactive microglia showed a smaller increase than in CTRL, mostly caudal to the injury site in the grey matter, indicating an early reduction of the inflammatory response attributable to cAMP delivery.

Protoplasmic and fibrous astrocytes were oppositely regulated, showing a significant reduction and increase respectively, in all injured conditions and along all spinal cord areas.

Pericytes and vascular endothelial cells were strongly decreased across all ROIs, segments, conditions and timepoints, with the decrease in caudal ROIs attenuated for both CTRL and STIM, most notably at 30 dpi in the dorsal white matter for vascular endothelial cells.

MOL and NFOL showed broad, sustained decreases with no recovery by 30 dpi, indicating persistent loss of mature and newly formed oligodendrocytes, although in the caudal ventral grey matter STIM samples showed lower MOL abundance than CTRL at both timepoints.

Against this background, myelinating oligodendrocytes (MFOL) and peripheral immune cells were the two populations most clearly restructured by stimulation. MFOL was strongly increased in STIM at both timepoints in every ROI of both segments, while CTRL moved in the opposite direction in most ROIs of the caudal segment, decreasing at both timepoints, or showed a smaller increase than STIM in most ROIs of the dorsal segment, consistent with a stimulation-driven acceleration of an early myelination-associated response. Peripheral immune cells were decreased in CTRL at both timepoints and in all ROIs, whereas STIM remained at or slightly above baseline at both timepoints, with no rostro-caudal gradient, implying persistent infiltration of immune cells under stimulation even at later stages.

Neuronal subtypes were detected only where expected (DE Sox5⁺Tac1⁺ and DE Reln⁺ dorsally, motor neurons and V1 interneurons medially and ventrally, VEP and MI caudally), and shared a common shape: injury-associated decreases that were attenuated in STIM at 10 dpi. By 30 dpi, the attenuation was largely lost in some cases: caudally, V1 fell further in STIM than in CTRL, and VEP decreased markedly more in STIM in the caudal medial ROI. MI was the clearest exception, remaining stable in STIM while CTRL declined further.

To follow how individual subtypes evolve over the course of injury and stimulation, we used maSigPro [4,5] (see Methods) to test for abundance changes and identify subtypes differing significantly between CTRL and STIM at any timepoint, and examined CLR-normalized trajectories from SHAM through 10 and 30 dpi for the significant neuronal subtypes (Fig. 3C).

Rostrally, DE Sox5⁺Tac1⁺ and motor neurons both fell from SHAM levels by 10 days in CTRL animals, and the initial drop was smaller under STIM. By 30 days, however, DE Sox5⁺Tac1⁺ converged on similarly reduced levels, while motor neurons showed some recovery in the STIM group. Rostral V1 interneurons behaved differently: CTRL declined less than STIM at 10 days, after which STIM partially recovered while CTRL continued to fall, indicating a possible recovery of V1 abundance at chronic stages.

Caudally we observed two patterns. First, V1 interneurons in the medial and ventral ROIs both dropped sharply in CTRL at 10 days and recovered partially by 30 days, whereas STIM showed only a modest decrease at 10 days; that early protection was transient, and both populations converged on CTRL levels by 30 days. Second, DE Reln⁺ and VCL V0d V2b showed opposite effects: the former was held at SHAM levels under STIM with a drop in CTRL, while for the latter the loss occurred in STIM samples instead.

bPAC-mediated stimulation therefore acts primarily on the early phase of injury-associated neuronal loss. Dorsal excitatory subtypes and motor neurons are buffered against the 10-day decline, most strongly caudal to the injury epicenter. This protection is not sustained, however, and most populations converge on CTRL levels by 30 days. V1 interneurons are the exception in both direction and ROI: they are spared early only caudally, and decline below CTRL by 30 days.

To further evaluate the spatial localization and condition-dependent abundance of the cell types inferred through deconvolution, we performed immunohistochemical analyses on coronal spinal cord sections with consensus cell markers for glial populations (Fig. 3D, left panels), neurons and blood vessels (Fig. 3D, right panels) in all conditions at 10 dpi. Olig2 nuclear expression (green) evidenced its dominant abundance alongside the oligodendroglial lineages in white and grey matter, as defined in Fig. 2E, and MBP staining (red) outlined the myelinated white matter. GFAP staining (green) identified astrocytes, showing stronger staining in the CTRL sample and indicating the increased number of fibrous astrocytes previously shown in the deconvolved spatial transcriptomics analysis (Fig. 3B). Microglia and infiltrated macrophages were identified by Iba1 immunostaining (green), which revealed sparse, low positive cell abundance in the SHAM sample compared with the dense accumulations of positive cells in the CTRL case. The STIM condition showed an Iba1 pattern comparable to SHAM, confirming the spatial transcriptomics analysis for this timepoint.

Blood vessel network density and architecture were visualized by CD31 (PECAM-1) endothelial-specific staining (red), outlining and confirming the transcriptomic analysis of an attenuated and disorganized blood vessel distribution in the injured samples, CTRL and STIM, in comparison with SHAM. Grey matter neuronal preservation and distribution were evaluated by NeuN (Rbfox3) staining (magenta). The reduced NeuN immunoreactivity in the CTRL sample compared to SHAM was prevented in the STIM condition, again confirming the transcriptional analysis, with a global neuronal preservation balance taking all neuronal cell populations into consideration. EN1 staining (green) marked V1 interneurons and ChAT staining (red) marked cholinergic motor neurons, the two populations whose deconvolved abundance was most clearly modulated by stimulation.

### Priority ranking of neuronal-function genes exposes an anatomically distinct transcriptional response

To identify genes whose expression dynamics may contribute to neuronal function during the injury and stimulation time course, we applied maSigPro [4,5] to the spatial transcriptomics dataset. The pipeline proceeded in five steps (Fig. 4A). First, Visium spots were summed into pseudobulk profiles per grey-matter area and section (dorsal, medial and ventral grey matter; three biological replicates per group). Second, genes were filtered and normalized with filterByExpr [30] and TMM [31], retaining those belonging to curated gene lists of spinal cord injury processes. Third, the resulting expression trajectories were modelled with maSigPro (negative-binomial GLM, time degree 2, Q < 0.05, R² > 0.4), isolating the genes whose temporal behaviour is significantly shaped by injury and by stimulation.

**Figure 4.**
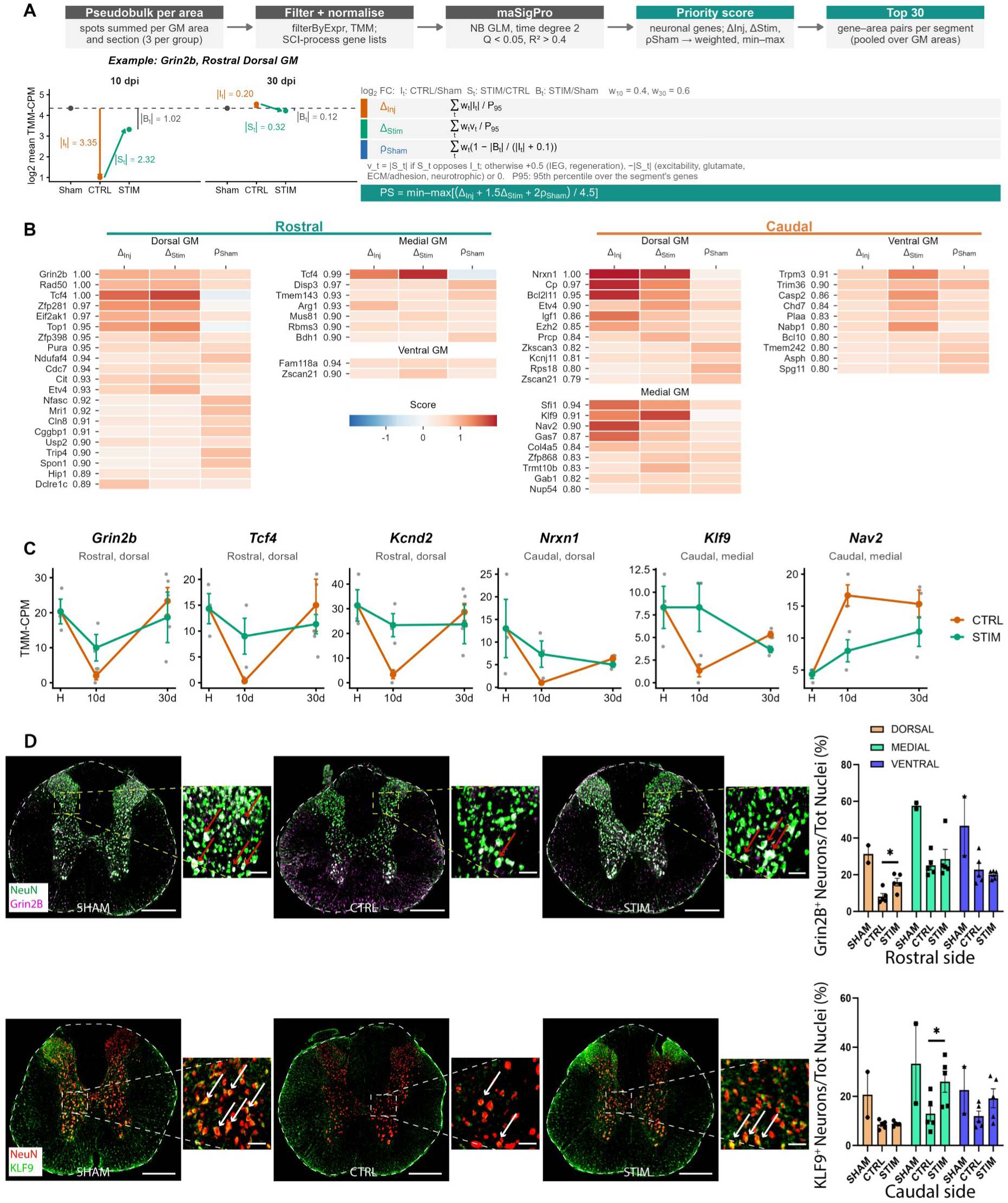
Spatial transcriptomic dynamics of neuronal-function genes following spinal cord injury and bPAC-mediated stimulation. **A,** Analysis pipeline: pseudobulk aggregation of spots per grey-matter area and section (three per group); filtering and normalization with filterByExpr and TMM, restricted to curated spinal cord injury process gene lists; maSigPro modelling (negative-binomial GLM, time degree 2, Q < 0.05, R² > 0.4); priority scoring of neuronal genes from ΔInj, ΔStim and ρSham, weighted and min-max scaled; and retention of the top 30 gene-area pairs per segment, pooled over grey-matter areas. Below, the worked example of Grin2b in rostral dorsal grey matter, showing the log2 mean TMM-CPM at 10 and 30 dpi with the injury, stimulation and baseline-residual magnitudes, alongside the definitions of the three components and of the final priority score. **B,** Priority-score heatmaps for the top-ranked genes of each segment, split by grey matter area: rostral (dorsal, medial, ventral) and caudal (dorsal, medial, ventral). Each gene is shown with its three components, ΔInj, ΔStim and ρSham, on a diverging colour scale, and with its final priority score beside the gene name. Components are normalized within each segment, so scores are comparable between areas within a segment but not between rostral and caudal. **C,** TMM-CPM expression across SHAM (H), 10 and 30 days for Grin2b, Tcf4 and Kcnd2 in rostral dorsal grey matter, Nrxn1 in caudal dorsal grey matter, and Klf9 and Nav2 in caudal medial grey matter. Data are mean ± SEM with individual replicates overlaid; CTRL in orange, STIM in green. **D,** Immunofluorescence and quantification at 10 dpi. Upper, NeuN (green) and Grin2B (magenta) in SHAM, CTRL and STIM sections, with boxed regions shown magnified and arrows marking double-positive neurons; right, quantification of Grin2B⁺ neurons as a percentage of total nuclei in dorsal, medial and ventral grey matter on the rostral side. Lower, NeuN (red) and KLF9 (green) with the equivalent magnifications and quantification on the caudal side. Asterisks denote a significant difference between CTRL and STIM. Scale bars as indicated.

These six analyses identified 514 significantly differentially expressed genes in total, with more than twice as many recovered caudally (350) as rostrally (164; Supplementary Table S1). The caudal excess was not explained by differences in sampling depth: in the medial grey matter, caudal samples contributed fewer spots and lower sequencing depth than rostral samples (36.9 vs 51.6 spots per sample; 1.13 × 10⁶ vs 1.26 × 10⁶ mean UMI) yet yielded more significant genes (63 vs 42). The asymmetry was most pronounced ventrally, where a 1.2-fold difference in spot number accompanied a 3.9-fold difference in gene recovery (140 vs 36), giving the highest normalized yield of any ROI, at 20.5 significant genes per 100 spots and 2.1% of all tested genes. The caudal dorsal grey matter produced the largest absolute number of genes (147), though it was also the most deeply sampled ROI, so this count is less readily separable from statistical power. Together, these results point to more extensive stimulation-dependent transcriptional remodelling caudal to the injury epicenter, consistent with the caudal preservation of neuronal abundance described above.

To prioritize genes for downstream interpretation, we first retained only genes expressed in at least 1% of neurons and with a mean expression of at least 0.02 in the single-cell data. We then ranked these genes using a custom priority score (Fig. 4A). For each retained gene, three quantitative descriptors were computed from the log2 fold-changes at 10 and 30 days, using time weights of 0.4 and 0.6 respectively to privilege the more chronic phase. ΔInj is the weighted magnitude of the injury effect (CTRL versus SHAM), normalized to the 95th percentile of the segment. ΔStim is the weighted magnitude of the stimulation effect (STIM versus CTRL), counted only when it opposes the injury effect; otherwise the gene process class determines the value, with immediate-early and regeneration genes receiving +0.5 and excitability, glutamate, extracellular matrix, adhesion and neurotrophic genes penalized. ρSham is the proximity of the stimulated condition to the SHAM baseline, quantified as the residual distance of STIM from SHAM relative to the injury displacement. The three descriptors were combined into the priority score, which assigns the greatest weight to the completeness of the return to the SHAM state, followed by the magnitude of the opposing stimulation effect and of the injury effect itself; the detailed parameterization is described in the Methods.

The pipeline is illustrated in Fig. 4A by a worked example, Grin2b in the rostral dorsal grey matter: the injury collapses its expression at 10 days (absolute injury effect 3.35), stimulation restores it (absolute stimulation effect 2.32), and by 30 days the stimulated condition lies close to the SHAM baseline (residual 0.12), yielding ΔInj = 1.02, ΔStim = 0.92, ρSham = 0.63 and the maximal priority score of 1.00. Genes were then ranked by priority score and the top 30 gene-area pairs per segment, pooled over the grey-matter areas, were retained. These top-ranked genes constitute the candidate effectors of the mechanism of action of bPAC treatment.

Figure 4B shows the top 30 priority genes for each spinal cord section as row-scaled expression heatmaps, together with the contribution of each component to the final priority score. The distribution of high-priority genes differed markedly between sections. In the caudal section, these genes were distributed relatively evenly across the dorsal (11 genes), medial (9 genes) and ventral (10 genes) areas, whereas the rostral section showed a pronounced enrichment in the dorsal area (21 genes). These findings suggest that the transcriptional response of neuronal-function genes to injury and stimulation is anatomically distinct between rostral and caudal spinal cord sections. We further observed that, with the exception of Tcf4 in the medial area, most genes received higher priority scores in the caudal than in the rostral section. This is consistent with the greater number of genes identified along the caudal section and suggests that the caudal cord exhibits a more extensive transcriptional response to injury and stimulation. One possible explanation is that the caudal samples capture the response of tissue downstream of the injury, where neuronal connections and their associated signalling pathways may be affected by the lesion. This anatomical distinction may therefore provide insight into how injury and stimulation responses are distributed along the spinal cord.

Furthermore, the three components of the priority score separate the top-ranked genes of Fig. 4B into two groups that behave in opposite ways. The first combines a high injury component with a high stimulation component and a low, in some cases negative, baseline-return component: genes strongly displaced by the injury and strongly responsive to stimulation that are nevertheless still far from SHAM levels at 30 days. Tcf4, Zfp281 and Top1 in the rostral dorsal and medial areas are the clearest examples, and Nrxn1, Cp, Bcl2l11, Igf1 and Ezh2 occupy the equivalent positions caudally. The second group inverts the profile, with modest injury and stimulation components alongside the highest baseline-return values of their panel. Nfasc, Mri1, Cln8, Cggbp1, Trip4, Spon1 and Hip1 rostrally, and Zkscan3, Kcnj11, Rps18, Tmem242, Asph and Spg11 caudally, are deflected less by the lesion but are returned to uninjured levels by 30 days under stimulation.

The two groups also differ in the kind of genes they contain. The high-magnitude, poorly recovering group is dominated by upstream regulatory machinery: transcriptional and epigenetic regulators (Tcf4, Zfp281, Klf9, Ezh2), growth-factor and apoptotic signalling (Igf1, Bcl2l11), cytoskeletal and axon-growth effectors (Nav2, Gas7) and synaptic adhesion molecules (Nrxn1). These are the components that would need to be redeployed to rebuild circuitry, and they remain persistently reset a month after injury. The recovery group is instead weighted towards homeostatic maintenance: bioenergetic and metabolic enzymes (Bdh1, Mri1, Tmem242, Kcnj11), ribosomal and protein-turnover transcripts (Rps18, Zkscan3, Hip1), and the structural machinery of the axo-glial interface (Nfasc, Spon1, Spg11).

### Stimulation buffers opposite gene-expression responses in the rostral and caudal cord

To examine the highest-confidence hits in greater detail, we plotted TMM-normalized expression trajectories across the time course for selected high-priority genes in the rostral dorsal and the caudal dorsal and medial grey matter (Fig. 4C). These trajectories showed a clear anatomical difference in the direction of the injury response, whereas stimulation mainly attenuated the injury-associated change.

In the rostral dorsal grey matter, all three genes (Grin2b, Tcf4 and Kcnd2) dropped sharply at 10 dpi in CTRL animals, to near-undetectable levels for Tcf4. The reduction was substantially smaller under STIM. Grin2b expression fell to about 10% of SHAM levels in CTRL animals but stayed at about half of SHAM under STIM. Tcf4 and Kcnd2 kept about 60% and 75% of SHAM expression under STIM, respectively.

Grin2b staining supported the mRNA differential expression, showing significant expression recovery in the dorsal horn compared with the CTRL condition at 10 dpi, though without reaching SHAM levels (Fig. 4D, upper panels). This pattern suggests that bPAC stimulation attenuates the early suppression of genes involved in synaptic signalling, neuronal excitability and transcriptional regulation.

By 30 dpi, CTRL expression had rebounded to approximately SHAM levels for all three genes, with a modest overshoot for Grin2b. STIM expression was closer to SHAM for Grin2b and Kcnd2 than for Tcf4, which stayed at about 80% of its SHAM level. Stimulation therefore mainly limited the depth of the early loss rather than changing the final 30 dpi level. Tcf4 was also the only gene in this group with a negative baseline-return score (Fig. 4B), consistent with its incomplete return to the uninjured state despite its maximal overall priority score.

The caudal cord showed a contrasting and more heterogeneous pattern. In the medial grey matter, Nav2 showed sustained induction rather than early suppression. In CTRL animals, its expression rose almost four-fold by 10 dpi and stayed elevated at 30 dpi. Under STIM, the rise was much smaller and more gradual, so expression remained below CTRL levels at both timepoints. This suggests that stimulation dampens a sustained injury-associated response involving cytoskeletal and neuronal growth processes.

In the caudal dorsal grey matter, Nrxn1 decreased sharply at 10 dpi, with a less severe reduction under STIM. By 30 dpi, however, both groups had converged on levels well below SHAM. Nrxn1, a key component of synaptic adhesion and neuronal connectivity, therefore remained jointly depressed rather than recovering to the uninjured state. This fits its near-zero baseline-return score despite its maximal overall priority score (Fig. 4B).

Klf9, in the caudal medial grey matter, followed a pattern similar to the rostral genes at 10 dpi. Injury strongly reduced its expression in CTRL animals, whereas STIM animals kept SHAM-like levels. Staining confirmed this effect: at 10 dpi, representative spinal cord immunostaining images and quantification of double-positive cells (NeuN⁺/KLF9⁺; Fig. 4D, lower panels) showed that the injury-dependent loss of KLF9⁺ neurons in the medial grey matter was significantly reversed by stimulation. By 30 dpi, however, Klf9 expression was below SHAM in both groups, and lower in STIM than in CTRL animals. The early protective effect of stimulation on Klf9 was therefore not maintained at the transcript level.

Overall, the rostral response was characterized by an early, stimulation-attenuated loss of expression followed by a rebound. The caudal response was more heterogeneous, including sustained injury-driven upregulation partially counteracted by stimulation (Nav2), persistent joint depression (Nrxn1), and transient preservation of expression under stimulation (Klf9). This regional difference is consistent with the distribution of the top 30 priority genes across the spinal cord sections (Fig. 4B). The trajectories also show that a high priority score does not necessarily indicate recovery towards the SHAM condition: genes can be strongly affected by injury and treatment and still remain transcriptionally displaced at 30 dpi. Together, these findings suggest that bPAC stimulation modulates the temporal dynamics of neuronal-function-associated genes in a regionally distinct manner. Its clearest effect is on the magnitude of the early injury response, with more variable effects on the persistence and resolution of transcriptional changes.

## Discussion

### Spatial and temporal transcriptomic architecture of bPAC-mediated recovery

Light-driven adenylyl cyclase (bPAC) activation offers precise spatiotemporal control over intracellular cyclic adenosine monophosphate (cAMP) dynamics [2]. Following spinal cord injury, endogenous cAMP levels decline within the lesion microenvironment, impairing intrinsic axonal growth capacity and exacerbating secondary damage cascades.

Our spatial transcriptomic profiling across rostral and caudal spinal segments demonstrates that targeted bPAC optogenetic photostimulation reprograms the post-SCI transcriptional landscape. Rather than inducing global transcriptional shifts, bPAC elicits ROI-specific transcriptional responses, orchestrating early structural matrix remodelling and neuroprotective pathways in rostral segments while driving remyelination programmes, axonal maintenance and immune suppression in caudal tissue.

### PKA/CREB-mediated neurotrophic priming and axonal regeneration

A central downstream effector of bPAC-induced cAMP elevation is protein kinase A (PKA), which phosphorylates the cAMP response element-binding protein (CREB). Phosphorylated CREB drives the transcription of pro-regenerative genes that enable axons to overcome extrinsic myelin-associated inhibitors.

In rostral grey matter, bPAC stimulation drives a late-phase upregulation of insulin-like growth factor 2 (Igf2), reaching robust expression levels at day 30 post-SCI, exceeding 1,400 normalized counts in bPAC samples compared to controls. IGF2 signalling acts synergistically with cAMP/PKA to activate downstream PI3K-Akt and mTOR pathways, promoting motoneuron survival and neurite elongation. Concurrently, early neuropeptide responses such as galanin (Gal) and urotensin 2 (Uts2) are markedly induced by bPAC. Galanin is a CREB-responsive neuropeptide that enhances axon sprouting, suggesting that bPAC reactivates an intrinsic embryonic-like growth programme within surviving spinal circuits.

bPAC treatment also modulates key transcriptional regulators, including activating transcription factor 4 (Atf4) and capicua (Cic). Atf4, a bZIP transcription factor downstream of cAMP/PKA and the integrated stress response, exhibits sustained, balanced expression in bPAC-treated rostral segments. This selective activation avoids chronic pro-apoptotic signalling, such as CHOP induction, and instead favours adaptive protein folding, antioxidant defence and metabolic resilience under ischaemic stress.

### EPAC/Rap1 signalling, node architecture and cytoskeletal stabilization

Beyond PKA, cAMP directly engages exchange proteins directly activated by cAMP (EPAC1/2). EPAC signalling activates small GTPases such as Rap1 and Rac1, governing cell adhesion, microvascular integrity and cytoskeletal dynamics independently of transcription factor phosphorylation.

The structural integrity of nodes of Ranvier is vital for saltatory conduction along injured long tracts. Neurofascin (Nfasc), a cell-adhesion molecule required for nodal clustering, exhibits severe dysregulation following unstimulated SCI but is stabilized under bPAC stimulation across both rostral and caudal segments. Concurrently, the structural neuronal markers ankyrin 2 (Ank2) and microtubule-associated protein 2 (Map2) remain robustly expressed in caudal segments under bPAC treatment. EPAC/Rap1 activation promotes the anchoring of NFASC and ANK2 to the actin-spectrin cytoskeleton, which would prevent focal demyelination and nodal disassembly caudal to the lesion.

Tenascin C (Tnc) and periostin (Postn) were induced by injury in rostral grey matter by 10 days in untreated animals. Stimulation did not enhance this early response: Tnc was lower under bPAC than in injured controls at 10 days in all three rostral areas, as was Postn in dorsal grey matter, whereas by 30 days Tnc was higher under bPAC in dorsal and medial grey matter, and Postn in dorsal grey matter. Stimulation therefore appears to delay rather than accelerate matrix gene induction. Because acute matrix deposition supports lesion stabilization while persistent deposition contributes to fibrotic scarring, the functional consequence of this shifted profile cannot be inferred from transcript levels and would require histological assessment of the lesion boundary.

### Preservation of oligodendrocyte bioenergetics and caudal remyelination

Secondary injury after SCI leads to oligodendrocyte loss, demyelination and subsequent functional conduction block [13]. cAMP signalling is a master regulator of oligodendrocyte progenitor cell differentiation and mitochondrial bioenergetics.

In caudal spinal segments, unstimulated SCI leads to a prolonged decline in major myelin structural transcripts. By contrast, at 30 days these transcripts were expressed at higher levels under bPAC stimulation than in untreated animals, although most were lower at 10 days. Elevated cAMP drives PKA phosphorylation of myelin gene promoters, while SIRT2 maintains oligodendrocyte metabolic homeostasis. This signature is consistent with preserved oligodendrocyte function under stimulation, which histological assessment of myelination would be needed to confirm.

Optogenetic elevation of cAMP also regulates cellular metabolism to meet the bioenergetic demand of axonal repair and remyelination. High-output metabolic genes, including creatine kinase B (Ckb), phosphoglycerate mutase 1 (Pgam1) and glycerol-3-phosphate dehydrogenase 1 (Gpd1), show preserved, elevated counts in bPAC-treated tissue. By restoring ATP generation through creatine-phosphate shuttling and glycolytic flux, bPAC would protect neurons and glia from energy collapse during the acute post-traumatic phase.

### Neuroimmune resolution and microglial modulation

Persistent neuroinflammation driven by activated microglia and infiltrating macrophages drives secondary tissue loss after SCI [13]. Elevated intracellular cAMP is a canonical anti-inflammatory signal in myeloid cells.

Unstimulated caudal segments show prolonged, high-level expression of neuroinflammatory transcripts, including MHC class II molecules (RT1-Bb, RT1-Db1) and lysozyme 2 (Lyz2), while chemokine ligand 5 (Ccl5) is modulated rostrally, bPAC stimulation lowered these transcripts at 30 days. Because cAMP/PKA signalling is known to inhibit NF-κB transcriptional activity by preventing p65 nuclear translocation, this mechanism could account for the effect.

### Conclusion and model of bPAC action

In summary, our transcriptomic data support a model in which bPAC photostimulation functions as a multi-modal molecular catalyst. In the acute phase, at day 10, bPAC attenuates the early induction of matrix genes (Tnc, Postn) rostrally while preserving bioenergetic pathways (Ckb, Pgam1) to limit acute cell death. In the chronic recovery phase, at day 30, bPAC sustains cAMP/PKA/CREB neurotrophic cascades (Igf2, Gal) rostrally, while resolving caudal neuroinflammation (RT1-Bb, Lyz2) and reactivating oligodendrocyte myelin repair machinery (Plp1, Mag, Cnp).

By targeting multiple downstream cAMP effectors simultaneously across time and space, bPAC optogenetic stimulation transforms the inhibitory post-SCI microenvironment into a permissive domain for neuroprotection, axonal maintenance and structural repair.

The pathway mechanisms proposed here, namely PKA/CREB, EPAC/Rap1 and NF-κB signalling, are inferred from the established biology of cAMP rather than measured directly.

## Methods

### Rat model and experimental design

Fifteen female Sprague Dawley rats of about 2 months of age (200 g) were distributed into 5 groups of 3 animals each: Sham, SCI 10, SCI 30, STIM 10 and STIM 30. Sham and SCI groups received intracortical injections of AAV9 CAMK2-bPAC-L123R-mCherry, the inactive mutant, while AAV9 CAMK2-bPAC-mCherry was injected in the STIM groups. Stereotaxic injections were performed to target layer V of the motor cortex at the hindlimb cortical areas as previously described, with 10 injection sites of 2 µL of 1 × 10¹³ vg/mL per hemisphere, using the PV850 microinjector (World Precision Instruments, USA). A bilateral brain lead device was then implanted in all rats to allow blue light stimulation at 470 nm (Neurolux, Northfield, USA) at the injection site, as described by Martínez-Rojas and colleagues [1]. All animals were subjected to light stimulation from day 1, once a day for 30 minutes until sacrifice. During stimulation, animals were placed in a cage connected to a power distribution control box delivering wireless power to the implants, allowing non-conditioned locomotion analysis during stimulation. The efficiency of blue light for cortical stimulation was confirmed by the increased amount of cAMP in the stimulated cortical area [1].

All surgical procedures were performed under anaesthesia, analgesia and antimicrobial treatment: 2% isoflurane in a continuous oxygen flow of 1 L/min, a single subcutaneous dose of morphine (2.5 mg/kg) for the surgical procedure, post-surgical buprenorphine (1 mg/kg every 8 hours for 4 days) and enrofloxacin (5 mg/kg every 24 hours for 7 days).

Sham, SCI 30 and STIM 30 underwent the surgical procedure for laminectomy and SCI 20 days before SCI 10 and STIM 10, and cortical injections were performed one week earlier accordingly. All animals were sacrificed on the same day by lethal dose of pentobarbital (100 mg/kg), followed by intracardiac perfusion with 50 mL of cold 0.9% NaCl solution to wash blood from the circulatory system. Spinal cords were dissected immediately after perfusion. The spinal cord was sectioned at the injury epicenter, or the equivalent level for the Sham group, and 0.5 cm rostral and 0.5 cm caudal segments were designated for spatial transcriptomics and immunostaining. The 0.5 cm segment immediately rostral was designated for single-nucleus isolation and snRNA-seq library preparation, as indicated in Fig. 1A. Each fragment was immersed in OCT matrix and immediately frozen using liquid nitrogen and isopentane, strictly following Visium and 10x Genomics instructions for spatial transcriptomics analysis to avoid RNA degradation. Samples were stored at −80 °C before processing.

### snRNA-seq library preparation

Nuclei were isolated using a protocol adapted from Matson and colleagues [22], including meninges removal, mechanical dissociation and sucrose-based differential centrifugation, followed by counting and resuspension. We obtained between 600 and 3,000 nuclei/µL across animals and collected an equal number of nuclei (60,000) from each rat to build five pooled samples, one for each experimental condition. All pools were adjusted to 1,500 nuclei/µL and processed following the Chromium Gem-X 10x Genomics single-cell protocol [23]. cDNA quality control was satisfactory, yielding 40 µL of cDNA per pool, of which 10 µL were used for Illumina library preparation. Post-library quality control confirmed adequate fragment size distributions and concentrations before sequencing by Macrogen.

### Spatial transcriptomics library preparation

Frozen tissues were cryosectioned immediately rostral and caudal to the injury epicenter, or the equivalent level for the Sham group. The first 20 µm sections per sample were collected when the injury-induced cavity was not detected, and mounted on Visium Gene Expression slides [20], yielding 30 spatial transcriptomics samples: 15 rostral and 15 caudal sections, randomized and distributed across several Visium slides. H&E staining and imaging confirmed good tissue morphology before permeabilization. After cDNA amplification, strictly following the manufacturer instructions, we obtained 40 µL of cDNA per spatial sample for library generation, submitted to Macrogen for sequencing.

### Read alignment

Raw FASTQ files from 10x Chromium single-nucleus RNA sequencing were processed using Cell Ranger v8.0.1 [23]. A custom Rattus norvegicus reference genome was first constructed using spaceranger mkref, based on the Rnor_6.0 genome assembly (GCF_000001895.5, NCBI RefSeq) and its corresponding GTF annotation file. Count matrices were generated from the raw output for each of the 5 single-nucleus samples. Prior to quality control, ambient RNA contamination was removed from raw count matrices using CellBender v0.3.0 [24] with remove-background under default parameters, using the raw unfiltered feature-barcode matrix as input. CellBender-corrected filtered count matrices were then used as input for all downstream preprocessing and quality control steps.

### snRNA-seq preprocessing and quality control

Initial preprocessing and quality control of the snRNA-seq data were performed with Scanpy [25] using a data-driven adaptive filtering approach. To account for sample-specific technical variance, adaptive thresholds were computed using the median absolute deviation (MAD) from the sample median. Nuclei were flagged as structural outliers and removed if their log-transformed total counts or log-transformed number of detected genes deviated by more than 5 MADs from the median. Additionally, nuclei exceeding 3 MADs for ribosomal or mitochondrial transcript fractions, with a strict ceiling constraint of mitochondrial fraction below 5%, were discarded. Genes not expressed in at least one valid nucleus were dropped.

Following adaptive QC filtering, baseline doublet detection was performed on the raw count matrices using scDblFinder [26] under default parameters. To establish an ultra-pure reference dataset suitable for downstream spatial deconvolution, a secondary, highly stringent filtering step was applied prior to signature generation, removing all nuclei with a residual scDblFinder score greater than 0.01. This strict cutoff removed an additional 23% of nuclei relative to the default scDblFinder baseline. Robustness validation verified that this stringent threshold introduced no structural or biological bias: relative neuronal subtype proportions remained stable (Supplementary Figure Not shown), and global gene expression profiles maintained a near-perfect linear relationship before and after filtering.

### snRNA-seq integration and cell-type annotation

Pilot and full-experiment datasets were concatenated, log-normalized and reduced to 50 principal components computed on all genes. Batch effects between the two experiments were corrected with Harmony [27] (harmonypy v0.0.10, default parameters), using experiment as the batch variable, because each sample corresponds to a single condition. A neighbourhood graph of 15 neighbours was built on the Harmony-corrected embedding. Cell types were annotated with CellTypist v1.6.3 [19] using custom logistic regression models trained on the Tabulae Paralytica mouse spinal cord atlas [17]. Training used nuclei from uninjured animals and from 7 and 14 days post-injury in the atlas time course. Reference and query counts were normalized to 10,000 counts per nucleus and log-transformed. Models were trained with CellTypist feature selection, which retrains the classifier on the genes with the highest coefficients. Genes were matched between mouse and rat by symbol, and 2,212 to 2,219 of 2,358 model features, that is 93.8% to 94.1%, were detected in the query.

Predictions were refined by majority voting: the query was over-clustered on its neighbourhood graph, and each over-cluster was assigned its most frequent predicted label. Major cell classes were annotated at the coarsest atlas level of 15 cell types, separately for each experiment with identically trained models. Re-annotating both experiments with a single saved model reproduced 94% to 95% of per-nucleus predictions. Disagreements were confined to the microglia-peripheral immune boundary, subsequently resolved by marker-based curation as described below, and to adjacent oligodendrocyte maturation stages. Adding 1-month and hemisection nuclei to the reference mainly shifted CellTypist peripheral immune calls to microglia, consistent with the marker-based reassignment.

Because the atlas annotation level used does not distinguish reactive microglia, these were identified in the pilot dataset as a Gpnmb-high Leiden sub-cluster [28] of microglia, at resolution 0.5 computed on highly variable genes, and transferred to the full experiment by kNN majority voting with k = 20 on the Harmony-corrected embedding. Cells initially annotated by CellTypist as peripheral immune cells that showed a reactive-microglial profile (Gpnmb⁺ Igf1⁺ Sall1⁺ Hexb⁺, F13a1⁻ Mrc1⁻) were reassigned to reactive microglia. Peripheral immune cells were defined as the macrophage cluster of each dataset Leiden clustering at resolution 0.8 (F13a1⁺ Mrc1⁺ Cd163⁺ Lyve1⁺ Sall1⁻), followed by manual curation. A small population of low-signal nuclei initially annotated as NK/T cells but lacking lymphocyte markers, n = 165 or 0.14% of nuclei, was retained within the microglial classes. In sham samples, cells in the reactive cluster, n = 61, showed only weak reactive marker expression and were merged with microglia. All automated annotations were inspected manually for expression of canonical markers. The final curated labels were used to build the cell-type references for RCTD deconvolution.

For neuronal subtype annotation, we subset neurons from the major-class annotation, that is nuclei labelled as dorsal or ventral neurons by majority voting, in both experiments and combined them into a single object. A CellTypist model was trained on neurons from the same Tabulae Paralytica atlas [17] timepoints (uninjured, 7 and 14 days post-injury; 18,140 nuclei) at annotation level 4, covering 19 dorsal, ventral, intermediate and cerebrospinal fluid-contacting neuron subtypes. Training and preprocessing followed the major-class model, including feature selection. The model was applied jointly to pilot and full-experiment neurons, so that all conditions were labelled by a single classifier, and predictions were refined by majority voting. Motor neurons, which are not represented at this level of the atlas, were annotated manually by inspecting key markers such as Chat within the ventral excitatory population. Nuclei assigned to the atlas neural stem cell class within the neuronal hierarchy were excluded, as the class lacked specific markers, co-expressed oligodendrocyte genes (Mbp, Plp1, Mobp, Mog, Bcas1) and had roughly two-fold higher UMI and gene counts than other neurons. It was therefore considered to consist of residual doublets not detected by scDblFinder. Subtype assignments were confirmed by inspecting canonical marker expression.

To maximize computational efficiency and minimize transcriptomic noise, the feature selection functionality in CellTypist was enabled to isolate the most informative, cell-type-specific marker genes. The resulting trained model weights for these genes were exported as a serialized configuration for downstream query deployment. Preprocessed query expression matrices were then annotated using this custom neuronal classifier. To mitigate stochastic gene dropout and technical artefacts inherent to single-nucleus transcriptomics, individual cell-by-cell predictions were refined using a neighbourhood-based correction by majority voting. This approach contextualized raw classification probabilities against a local k-nearest neighbour graph to assign robust, smoothed consensus cluster identities. Automatically annotated cell types and neuronal subtypes generated using CellTypist were then inspected manually to confirm that gene markers were expressed in accordance with the assigned subtype.

### Spatial transcriptomics preprocessing and quality control

Raw FASTQ files from 10x Visium spatial transcriptomics [20] were processed using Space Ranger v3.0.1 [23] with the same custom Rattus norvegicus Rnor_6.0 reference genome described above. Count matrices were generated from the raw output for each of the 30 spatial transcriptomics samples.

Tissue-covered spots were manually selected for each section prior to alignment using Loupe Browser, and the resulting spot selections were exported as JSON alignment files. Read alignment and count matrix generation were then performed independently for each of the 30 spatial samples using spaceranger count, supplying the custom Rnor_6.0 reference, all FASTQ files across sequencing runs, the high-resolution brightfield tissue image, and the Loupe Browser alignment file for each sample. Slide identity and capture area were parsed automatically from image filenames. The filtered count matrices, spatial coordinates and tissue position files output by Space Ranger were used for all downstream analyses.

Quality control metrics, including total UMI counts, number of detected genes, and mitochondrial, ribosomal and haemoglobin transcript fractions, were computed per spot using Scanpy [25] and Squidpy [29] for inspection purposes. Outlier spots were flagged using MAD-based thresholds, 5 MADs for log-transformed total counts and gene counts and 3 MADs for mitochondrial and ribosomal fractions, and visualized spatially, but no automated spot filtering was applied at this stage, as tissue selection had already been performed manually in Loupe Browser.

### Alignment of spatial slices

To enable cross-sample spatial comparison, the spatial coordinates of all 30 sections were aligned into a common coordinate space using STIMA, Spatial Transcriptomics Image-based Methods for Alignment [3]. STIMA performs alignment using affine transformations applied simultaneously to the histological microscopy image and the gene expression spot matrix, relying exclusively on image-based information without incorporating transcriptomic data, thereby preserving the independence of gene expression across samples. Aligned spot coordinates were subsequently exported for use in all downstream regional analyses.

### Cell-type deconvolution

Spatial deconvolution of cell-type composition was performed using Robust Cell Type Decomposition (RCTD; spacexr v2.2.1) [21], run independently per sample in full mode, which allows each spot to be assigned weighted contributions from multiple cell types simultaneously. To maximize biological accuracy, each spatial sample was deconvolved against a condition-matched snRNA-seq reference built from pooled tissue of animals of the same experimental condition. This yielded three reference datasets: SHAM, CTRL and STIM. For the CTRL and STIM conditions, the references were built by merging both timepoints, 10 and 30 days post-injury, at the single-nucleus level. Each reference was constructed from the annotated count matrices, cell-type labels and per-cell total UMI counts of the corresponding snRNA-seq dataset described above, with a minimum cell instance threshold of 1. Following deconvolution, per-spot weight matrices of spots by cell types were extracted for each sample and used in all downstream cell-type proportion and differential abundance analyses.

### Differential cell-type abundance analysis

To assess condition-dependent changes in cell-type composition across spatially defined ROIs, we applied the maSigPro [4,5] regression framework to RCTD deconvolution weight matrices. Analyses were performed separately for rostral and caudal spinal cord sections, and for five manually delineated ROIs: dorsal, medial and ventral grey matter, and dorsal and ventral white matter.

For each sample, per-spot RCTD weights were normalized to sum to one. Entries below 5% of the spot total were set to zero, and each spot was re-normalized to sum to one. Weights were then summed per cell type across all spots of each ROI and divided by the total, giving the mean per-spot proportion of each cell type. This weights every spot equally and makes compositions independent of ROI size. Cell types with a mean proportion of 2% or less across samples within an ROI were removed, and the remaining proportions were re-closed to sum to one. Proportions were then transformed using the centred log-ratio, with a pseudocount of 10⁻⁴ added to all cell types to handle zeros.

CLR matrices were analysed with maSigPro. The design comprised SHAM, CTRL and STIM animals at 10 and 30 days post-injury, with n = 3 per group and timepoint and SHAM samples as the common time-zero reference of both groups. A second-degree polynomial design was built with make.design.matrix. Cell types with a significant regression model were first selected with p.vector using the Benjamini-Hochberg FDR [32] at Q < 0.05. Models were then refined by backward stepwise selection with T.fit at alpha = 0.05, and cell types with R² > 0.4 and significant STIM versus CTRL terms were extracted with get.siggenes. Deconvolution used the condition-matched references.

For visualization, log₂ fold-changes relative to SHAM were computed per cell type, ROI and group as the difference between the group mean CLR value and the SHAM mean CLR value, divided by the natural logarithm of 2. Cell types not retained in a given ROI, with mean proportion at or below 2%, are marked with a cross, and asterisks denote significant STIM versus CTRL trajectories identified by maSigPro. For neuronal cell types with a significant STIM versus CTRL trajectory, CLR abundances are shown over time as group means ± standard error with n = 3, and individual samples overlaid. SHAM samples are the common starting point of the CTRL and STIM trajectories.

### Differential gene expression profile analysis

To assess differences in gene expression profiles across conditions, we applied the maSigPro [4,5] methodology to our spatial transcriptomics data and experimental time-course design. For each grey-matter area, spot counts were first aggregated into pseudobulk profiles per sample. Lowly expressed genes were removed with filterByExpr from edgeR [30] using the condition-by-timepoint groups, and TMM normalization factors [31] were calculated on all retained genes. The maSigPro analysis was then restricted to genes belonging to the union of 15 curated gene lists representing spinal cord injury and neural function-related processes, compiled from Gene Ontology annotations, comprising 14,935 genes and provided as Supplementary Table S2. Library sizes and normalization factors were retained from the full filtered gene set, so that normalization reflected the whole expressed transcriptome rather than the restricted gene set. Between 6,336 and 8,058 genes were tested per area. TMM-normalized counts per million, rounded to integers, were used as input for the two-step regression approach implemented in maSigPro, in which non-flat genes were identified at Q < 0.05 and significant genes with R² > 0.4 retained, then clustered into temporal profiles.

### Gene priority score

To prioritize genes for downstream interpretation, we first retained only genes expressed in at least 1% of neurons and with a mean expression of at least 0.02 in the single-cell data. Scoring was restricted to these neuronal genes.

To facilitate biological interpretation of the maSigPro differential expression results, a composite priority score was developed to rank neuronal differentially expressed genes according to three biologically meaningful criteria, which constitute its three components: the size of their transcriptional response to spinal cord injury, how much bPAC stimulation counteracted that response, and how far stimulated animals returned toward the SHAM baseline.

For each ROI and segment, the expression profiles of the maSigPro-significant genes, as TMM-normalized CPM, were averaged across the three biological replicates of each condition. Log₂ fold changes were computed as the log₂ ratio of the corresponding group means for each timepoint: injured versus SHAM, stimulated versus injured, and stimulated versus SHAM, each at 10 and 30 days post-injury. Temporal contributions to the score components were weighted asymmetrically across the two post-injury timepoints, 0.4 at 10 days and 0.6 at 30 days, to emphasize the chronic phase, which better reflects the sustained transcriptional changes most relevant to therapy.

Because the rostral and caudal datasets were modelled independently, the score was computed separately for each segment. All normalization constants are therefore segment-specific, and priority score values rank genes within a segment rather than on a scale shared between them. Genes were additionally restricted to a curated set of transcripts with high neuronal expression prior to scoring. Within each segment, the 30 highest-scoring genes across all three grey matter subregions were retained for visualization; because selection was pooled rather than applied per subregion, the resulting panels contain unequal numbers of genes. The complete scored gene lists are provided as supplementary material.

### Component 1: injury magnitude (Δ_Inj)

The first component quantifies the magnitude of each gene’s transcriptional response to SCI, computed as the weighted absolute log₂ fold change between injured and sham conditions across both post-injury timepoints:

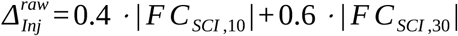

To prevent extreme outliers from dominating the ranking, raw scores were normalized by the 95th percentile of the population 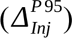:

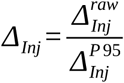

### Component 2: treatment effect (Δ_Stim)

The second component evaluates the rescue effect of bPAC stimulation. At each timepoint, the sign of the injury-induced fold change was compared with the sign of the bPAC-induced fold change. Where the two opposed, that is where stimulation moved expression counter to the injury effect, the timepoint value was set to the absolute magnitude of that rescue. Only where the signs did not oppose was the value instead determined by the gene functional category: genes in pro-regenerative categories, namely immediate-early and regeneration-associated genes, received a fixed positive value of 0.5; genes in categories associated with detrimental remodelling, namely excitability, glutamatergic signalling, extracellular matrix, matrix adhesion and neurotrophic, were penalized by the absolute magnitude of the bPAC effect; all remaining genes received 0. Component values across timepoints were aggregated and normalized by the 95th percentile of the population (*Δ^P^* ^95^, with a floor of 0.01):

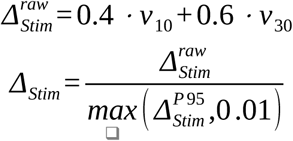

Consequently, the functional-category rules apply only to genes for which stimulation did not oppose the injury direction at that timepoint; for genes showing directional opposition, the category does not enter the calculation. Timepoint values were aggregated and normalised by the 95th percentile of the segment’s population, with a floor of 0.01 to guard against degenerate scaling:

This component may take negative values for genes penalised at both timepoints.

### Component 3: return to SHAM baseline (ρ_Sham)

The third component captures the extent to which bPAC stimulation restored expression toward healthy baseline levels. At each timepoint, the residual deviation of stimulated animals from sham, | *F C_StimvsSham, t_* |, was expressed relative to the injury-induced change (| *F C_SCI, t_* |+ 0.1) and subtracted from 1 so that higher values indicate closer restoration:

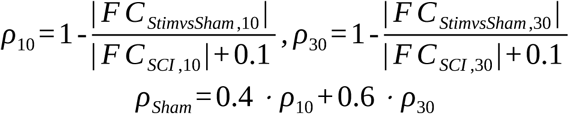

The constant 0.1 in the denominator is a regularisation term preventing division by zero for genes with negligible injury effects; it also limits the influence of such genes, whose ratios would otherwise be unstable. S_norm approaching 1 indicates near-complete restoration to healthy levels, while values near 0 indicate a residual deviation comparable to the injury effect itself. The component is unbounded below and takes negative values where the residual deviation exceeds the injury magnitude.

#### Final priority score

The three components were integrated using an explicit weighting scheme reflecting their relative biological importance: injury magnitude (W₁ = 1.0), treatment benefit (W₂ = 1.5), and return to healthy baseline (W₃ = 2.0, the highest weight, reflecting the primacy of functional recovery):

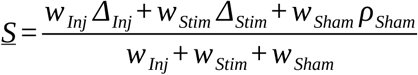

The weighted average was then rescaled to a [0, 1] interval using min-max normalization across all genes passing the maSigPro significance filters:

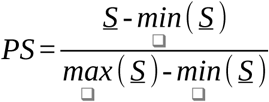

Because this rescaling is applied per segment, the highest-scoring gene in each of the rostral and caudal analyses takes a value of 1 by construction. PS therefore expresses a gene’s priority relative to the other genes in its own analysis and should not be compared in absolute terms between rostral and caudal samples. The same applies to S_SCI and S_bSCI, whose normalisation constants are segment-specific; S_norm contains no population-dependent term and is directly comparable across segments.

### Immunostaining

30 µm cryosections were collected in series immediately contiguous to the rostral and caudal slices designated for spatial transcriptomics for each sample. Standard immunofluorescence staining was performed for each series after short fixation with 4% paraformaldehyde in phosphate-buffered saline, followed by overnight incubation at 4 °C in primary antibody solution in TBS with 5% donkey serum and 0.25% Triton, and, after washing, one hour in secondary antibody solution. Primary antibodies were anti-GFAP (chicken, 1:500, PA1/10004, Invitrogen, USA), anti-choline acetyltransferase, ChAT (goat, 1:200, AB144P, Merck Millipore, Massachusetts, USA), anti-Olig2 (rabbit, 1:200, AB33427, Abcam, UK), anti-MBP (rabbit, 1:200, AB69863, Abcam, UK), anti-Iba1 (rabbit, 1:200, 019-19741, Fujifilm Wako, Japan), anti-NeuN (guinea pig, 1:400, ABN90, Millipore, USA), anti-EN1 (rabbit, 1:100, PA5-14149, ThermoFisher, Massachusetts, USA), anti-CD31 (goat, 1:50, AF3628, Biotechne, Minnesota, USA), anti-Grin2B (rabbit, 1:50, 718600, ThermoFisher, Massachusetts, USA) and anti-KLF9 (rabbit, 1:50, A7196, ABclonal, USA). Secondary antibodies were Alexa Fluor-488, -555 and -647 conjugated antibodies (1:400, Invitrogen, Massachusetts, USA). Cell nuclei were counterstained with DAPI (1:2000, Sigma, Missouri, USA). Sections were mounted in Mowiol and images for histological analysis were acquired on a Leica Thunder inverted microscope DMi8, equipped with a PL-APO 10×/1.40 air objective and a CMOS camera (Leica DFC9000 GTC). Köhler illumination was applied to enhance image contrast and to find the right plane with the bright-field channel. Excitation was achieved using the Spectra-X Light Engine LED lamp, which includes eight solid-state LEDs configured with Quad filter settings, enabling imaging of four different fluorophores without changing filters and significantly reducing acquisition time. The Leica Application Suite X software was used for microscope control and image acquisition, and image analysis and quantification were performed using ImageJ.

## Acknowledgements

The computations were performed on the HPC cluster Garnatxa at the Institute for Integrative Systems Biology (I2SysBio). I2SysBio is a mixed research centre formed by University of Valencia (UV) and Spanish National Research Council (CSIC).

## References

1. Martínez-Rojas, B., Martín-Pérez, S., Giraldo, E. et al. Stimulation of corticospinal neurons by optogenetic cAMP inductions promotes motor recovery after spinal cord injury in female rats via raphespinal tract modulation. Nat. Commun. 16, 5885 (2025). doi:10.1038/s41467-025-61018-3

2. Stierl, M., Stumpf, P., Udwari, D. et al. Light modulation of cellular cAMP by a small bacterial photoactivated adenylyl cyclase, bPAC, of the soil bacterium Beggiatoa. J. Biol. Chem. 286, 1181–1188 (2011). doi:10.1074/jbc.M110.185496

3. Gaya-Martín, V. A., Tarazona, S. & Conesa, A. STIMA: An R package for image-based spatial transcriptomics alignment (2026).

4. Conesa, A., Nueda, M. J., Ferrer, A. & Talón, M. maSigPro: a method to identify significantly differential expression profiles in time-course microarray experiments. Bioinformatics 22, 1096–1102 (2006). doi:10.1093/bioinformatics/btl056

5. Nueda, M. J., Tarazona, S. & Conesa, A. Next maSigPro: updating maSigPro bioconductor package for RNA-seq time series. Bioinformatics 30, 2598–2602 (2014). doi:10.1093/bioinformatics/btu333

6. Sathyamurthy, A., Johnson, K. R., Matson, K. J. E. et al. Massively parallel single nucleus transcriptional profiling defines spinal cord neurons and their activity during behavior. Cell Rep. 22, 2216–2225 (2018). doi:10.1016/j.celrep.2018.02.003

7. Häring, M., Zeisel, A., Hochgerner, H. et al. Neuronal atlas of the dorsal horn defines its architecture and links sensory input to transcriptional cell types. Nat. Neurosci. 21, 869–880 (2018). doi:10.1038/s41593-018-0141-1

8. Rosenberg, A. B., Roco, C. M., Muscat, R. A. et al. Single-cell profiling of the developing mouse brain and spinal cord with split-pool barcoding. Science 360, 176–182 (2018). doi:10.1126/science.aam8999

9. Zeisel, A., Hochgerner, H., Lönnerberg, P. et al. Molecular architecture of the mouse nervous system. Cell 174, 999–1014.e22 (2018). doi:10.1016/j.cell.2018.06.021

10. Blum, J. A., Klemm, S., Shadrach, J. L. et al. Single-cell transcriptomic analysis of the adult mouse spinal cord reveals molecular diversity of autonomic and skeletal motor neurons. Nat. Neurosci. 24, 572–583 (2021). doi:10.1038/s41593-020-00795-0

11. Russ, D. E., Patterson Cross, R. B., Li, L. et al. A harmonized atlas of mouse spinal cord cell types and their spatial organization. Nat. Commun. 12, 5722 (2021). doi:10.1038/s41467-021-25125-1

12. Alkaslasi, M. R., Piccus, Z. E., Hareendran, S. et al. Single nucleus RNA-sequencing defines unexpected diversity of cholinergic neuron types in the adult mouse spinal cord. Nat. Commun. 12, 2471 (2021). doi:10.1038/s41467-021-22691-2

13. Milich, L. M., Choi, J. S., Ryan, C. et al. Single-cell analysis of the cellular heterogeneity and interactions in the injured mouse spinal cord. J. Exp. Med. 218, e20210040 (2021). doi:10.1084/jem.20210040

14. Matson, K. J. E., Russ, D. E., Kathe, C. et al. Single cell atlas of spinal cord injury in mice reveals a pro-regenerative signature in spinocerebellar neurons. Nat. Commun. 13, 5628 (2022). doi:10.1038/s41467-022-33184-1

15. Yadav, A., Matson, K. J. E., Li, L. et al. A cellular taxonomy of the adult human spinal cord. Neuron 111, 328–344.e7 (2023). doi:10.1016/j.neuron.2023.01.007

16. Zhang, D., Chen, Y., Wei, Y. et al. Spatial transcriptomics and single-nucleus RNA sequencing reveal a transcriptomic atlas of adult human spinal cord. eLife 12, RP92046 (2024). doi:10.7554/eLife.92046

17. Skinnider, M. A., Gautier, M., Teo, A. Y. Y. et al. Single-cell and spatial atlases of spinal cord injury in the Tabulae Paralytica. Nature 631, 150–163 (2024). doi:10.1038/s41586-024-07504-y

18. Mejia, J. & Horner, P. J. Glia are bussin: how single-cell and spatial transcriptomics enlighten the role of neuroglia in spinal cord injury and regeneration. Glia 74, e70139 (2026). doi:10.1002/glia.70139

19. Domínguez Conde, C., Xu, C., Jarvis, L. B., et al. Cross-tissue immune cell analysis reveals tissue-specific features in humans. Science 376, eabl5197 (2022). doi:10.1126/science.abl5197

20. Ståhl, P. L., Salmén, F., Vickovic, S. et al. Visualization and analysis of gene expression in tissue sections by spatial transcriptomics. Science 353, 78–82 (2016). doi:10.1126/science.aaf2403

21. Cable, D. M., Murray, E., Zou, L. S. et al. Robust decomposition of cell type mixtures in spatial transcriptomics. Nat. Biotechnol. 40, 517–526 (2022). doi:10.1038/s41587-021-00830-w

22. Matson, K. J. E., Sathyamurthy, A., Johnson, K. R. et al. Isolation of adult spinal cord nuclei for massively parallel single-nucleus RNA sequencing. J. Vis. Exp. 140, e58413 (2018). doi:10.3791/58413

23. Zheng, G. X. Y., Terry, J. M., Belgrader, P. et al. Massively parallel digital transcriptional profiling of single cells. Nat. Commun. 8, 14049 (2017). doi:10.1038/ncomms14049

24. Fleming, S. J., Chaffin, M. D., Arduini, A. et al. Unsupervised removal of systematic background noise from droplet-based single-cell experiments using CellBender. Nat. Methods 20, 1323–1335 (2023). doi:10.1038/s41592-023-01943-7

25. Wolf, F. A., Angerer, P. & Theis, F. J. SCANPY: large-scale single-cell gene expression data analysis. Genome Biol. 19, 15 (2018). doi:10.1186/s13059-017-1382-0

26. Germain, P.-L., Lun, A., Macnair, W. & Robinson, M. D. Doublet identification in single-cell sequencing data using scDblFinder. F1000Research 10, 979 (2021). doi:10.12688/f1000research.73600.1

27. Korsunsky, I., Millard, N., Fan, J. et al. Fast, sensitive and accurate integration of single-cell data with Harmony. Nat. Methods 16, 1289–1296 (2019). doi:10.1038/s41592-019-0619-0

28. Traag, V. A., Waltman, L. & van Eck, N. J. From Louvain to Leiden: guaranteeing well-connected communities. Sci. Rep. 9, 5233 (2019). doi:10.1038/s41598-019-41695-z

29. Palla, G., Spitzer, H., Klein, M. et al. Squidpy: a scalable framework for spatial omics analysis. Nat. Methods 19, 171–178 (2022). doi:10.1038/s41592-021-01358-2

30. Robinson, M. D., McCarthy, D. J. & Smyth, G. K. edgeR: a Bioconductor package for differential expression analysis of digital gene expression data. Bioinformatics 26, 139–140 (2010). doi:10.1093/bioinformatics/btp616

31. Robinson, M. D. & Oshlack, A. A scaling normalization method for differential expression analysis of RNA-seq data. Genome Biol. 11, R25 (2010). doi:10.1186/gb-2010-11-3-r25

32. Benjamini, Y. & Hochberg, Y. Controlling the false discovery rate: a practical and powerful approach to multiple testing. J. R. Stat. Soc. B 57, 289–300 (1995). doi:10.1111/j.2517-6161.1995.tb02031.x

